# Identifying and exploiting trait-relevant tissues with multiple functional annotations in genome-wide association studies

**DOI:** 10.1101/242990

**Authors:** Xingjie Hao, Ping Zeng, Shujun Zhang, Xiang Zhou

## Abstract

Genome-wide association studies (GWASs) have identified many disease associated loci, the majority of which have unknown biological functions. Understanding the mechanism underlying trait associations requires identifying trait-relevant tissues and investigating associations in a trait-specific fashion. Here, we extend the widely used linear mixed model to incorporate multiple SNP functional annotations from omics studies with GWAS summary statistics to facilitate the identification of trait-relevant tissues, with which to further construct powerful association tests. Specifically, we rely on a generalized estimating equation based algorithm for parameter inference, a mixture modeling framework for trait-tissue relevance classification, and a weighted sequence kernel association test constructed based on the identified trait-relevant tissues for powerful association analysis. We refer to our analytic procedure as the Scalable Multiple Annotation integration for trait-Relevant Tissue identification and usage (SMART). With extensive simulations, we show how our method can make use of multiple complementary annotations to improve the accuracy for identifying trait-relevant tissues. In addition, our procedure allows us to make use of the inferred trait-relevant tissues, for the first time, to construct more powerful SNP set tests. We apply our method for an in-depth analysis of 43 traits from 28 GWASs using tissue-specific annotations in 105 tissues derived from ENCODE and Roadmap. Our results reveal new trait-tissue relevance, pinpoint important annotations that are informative of trait-tissue relationship, and illustrate how we can use the inferred trait-relevant tissues to construct more powerful association tests in the Wellcome trust case control consortium study.

**Author Summary:** Identifying trait-relevant tissues is an important step towards understanding disease etiology. Computational methods have been recently developed to integrate SNP functional annotations generated from omics studies to genome-wide association studies (GWASs) to infer trait-relevant tissues. However, two important questions remain to be answered. First, with the increasing number and types of functional annotations nowadays, how do we integrate multiple annotations jointly into GWASs in a trait-specific fashion to take advantage of the complementary information contained in these annotations to optimize the performance of trait-relevant tissue inference? Second, what to do with the inferred trait-relevant tissues? Here, we develop a new statistical method and software to make progress on both fronts. For the first question, we extend the commonly used linear mixed model, with new algorithms and inference strategies, to incorporate multiple annotations in a trait-specific fashion to improve trait-relevant tissue inference accuracy. For the second question, we rely on the close relationship between our proposed method and the widely-used sequence kernel association test, and use the inferred trait-relevant tissues, for the first time, to construct more powerful association tests. We illustrate the benefits of our method through extensive simulations and applications to a wide range of real data sets.

## Introduction

Genome-wide association studies (GWASs) have identified thousands of genetic loci associated with complex traits and common diseases. However, the majority (~90%) of these associated loci reside in noncoding regions and have unknown biological functions [1]. Systematic characterization of the biological function of genetic variants thus represents an important step for further investigating the molecular mechanisms underlying the identified disease associations. Functional characterization of genetic variants is challenging because the function and genetic effects of variants on most traits are likely acted through a tissue-specific fashion, despite their tissue-wide presence (certainly with the notable exception of somatic mutations) [2–4]. For example, it is well recognized that many psychiatric disorders, such as bipolar and schizophrenia, are consequences of dysfunctions of various genes, pathways as well as regulatory elements in neuronal and glia cells, resulting from brain-specific genetic effects of polymorphisms [5–9]. Therefore, characterizing the function of variants in various brain regions can help elucidate the biology of psychiatric disorders. For most complex traits, however, their trait-relevant tissues are often obscure. As a result, identifying trait-relevant tissues and characterizing the functions of genetic variants within relevant tissues holds the key for furthering our understanding of disease etiology and the genetic basis of phenotypic variation [10–16].

Both experimental and computational studies have recently produced a rich resource of variant annotations that can help characterize the function of genetic variants in a tissue-dependent fashion [17–21]. For example, the ENCODE and Roadmap epigenomics projects collect various measurements of histone modification, open chromatin, and methylation from more than a hundred different tissue and cell types, where each epigenetic mark characterizes a specific aspect of variant function [22,23]. Similarly, the GTEx project produces gene expression measurements from multiple tissues and quantifies variant function in terms of its ability to regulate gene expression levels in the given tissue [24]. Besides the experimental efforts, many computational methods have also been developed to create synthetic functional annotations for variants in a tissue-dependent manner. For example, the chromHMM converts measurements of multiple histone modifications in each tissue into 15 chromatin states that have more biologically interpretable functions than the original histone occupancy based annotations [25,26]. Similarly, several other methods provide different ways to summarize multiple SNP annotations into a single, potentially more interpretable annotation for various tissues [27,28].

With the large and growing number of tissue-specific variant annotations, however, one naturally wonders how these different annotations can be incorporated together to facilitate the identification of trait-relevant tissues. Several statistical methods have been recently developed to test the role of various functional annotations in predicting the variant effect sizes or causality for GWAS traits [10–16,29–31]. These methods often test one annotation at a time and produce a test statistic signifying the importance of the given annotation for a GWAS trait. By comparing the test statistics of univariate annotation from different tissues and ordering tissues by the magnitude of the test statistics, existing methods can be used to identify tissue relevance for a given trait [10,16]. However, examining one variant annotation at a time can be inefficient as it may fail to incorporate the rich information contained in various other annotations that likely characterize other aspects of variant function [27,28,32]. For example, some annotations are designed to evaluate evolutionary conservation of a variant, while some other annotations are designed to quantify its biochemical functionality [29,33]. Even annotations that belong to the same general category may characterize substantially different functions of a variant. For example, different histone modifications are used to annotate variants by different functional genomic regions: H3K4me3 annotates promoter; H3K4me1 annotates enhancer; H3K36me3 annotates transcribed regions; H3K27me3 annotates polycomb repression; H3K9me3 annotates heterochromatin; and H3K9ac annotates both enhancer and promoter [22,34]. Therefore, testing one annotation at a time may be suboptimal, and it would be ideal to incorporate multiple sources of information together to identify trait-relevant tissues. Besides the potential loss of inference efficiency, examining one annotation at a time can sometimes lead to incoherent results on the identification of trait-relevant tissues: partly due to a lack of statistical power, the trait-relevant tissues inferred by different SNP annotations may not always agree, and it is often not straightforward to consolidate results from using different annotations [27,28].

Despite the potential inefficiency due to the use of univariate annotation tests, several studies have explored the feasibility of inferring trait-relevant tissues for various complex traits using SNP functional annotations [16,31,35,36]. While the inferred trait-tissue relevance often makes biological sense, it is unclear how to further make use of these inferred trait-relevant tissues to benefit future association studies [11,16]. In principal, levering the information learned from the identified trait-relevant tissues could enable powerful association tests, as the functional annotations in the trait-relevant tissues could contain important SNP causality and effect size information. In practice, however, incorporating trait-relevant tissue information into association tests is challenging, partly because the existing statistical methods for identifying trait-relevant tissues mainly rely on polygenic models [16,31,35,36] while the existing statistical methods for association tests mostly rely on univariate tests or sparse regression models [37–40]. The disparity between the methods used for trait-tissue relevance inference and the methods used for association tests make it hard to share information across the two different tasks.

Here, we develop a simple method to address the above two challenges. First, we incorporate multiple binary and/or continuous annotations to facilitate the identification of trait-relevant tissues for GWAS traits. To do so, we modify the commonly used linear mixed model [38,41–45] to relate variant effect sizes to variant annotations by introducing variant specific variance components that are functions of multiple annotations. We quantify and evaluate the joint contribution of multiple annotations to genetic effect sizes by performing parameter inference using the widely used generalized estimation equation (GEE) [46]. Our GEE-based algorithm is closely related to the recent LDSC [16], ployGEE [47], and MQS methods [48], allows for the use of summary statistics, and naturally accounts for the correlation among summary statistics due to linkage disequilibrium. With GEE statistics, we further apply mixture models to classify tissues into two categories -- those that are relevant to the trait and those that are not -- thus formulating the task of identifying trait-relevant tissues into a classification problem. Second, our method is closely related to the sequence kernel association test (SKAT) [49–51], and this relationship allows us to apply parameter estimates from the inferred trait-relevant tissues as SNP weights to construct SNP set test and power new association studies. We refer to our overall analytic procedure as the Scalable Multiple Annotation integration for trait-Relevant Tissue identification (SMART). We provide an overview of our method in the Materials and Methods section, with details described in the Supplementary Text. With simulations, we show that, compared with analyzing one annotation at a time, analyzing multiple annotations jointly can improve power for the identification of trait-relevant tissues. In addition, we show that, using parameter estimates from inferred trait-relevant tissues as SNP weights leads to more powerful SNP set tests than the standard SKAT [49–51]. We apply our method for an in-depth analysis of 43 GWAS traits with multiple functional annotations in more than one hundred tissues derived from ENCODE and Roadmap. We show how our method and analysis can help provide biological insights for the genetic basis of complex traits and benefit future association studies. The SMART method is implemented as an R package, freely available at http://www.xzlab.org/software.html.

## Materials and Methods

### Method Overview

We consider a simple extension of the linear mixed model to evaluate jointly the contribution of multiple SNP annotations. To do so, we first consider the following multiple linear regression model that relates genotypes to phenotypes

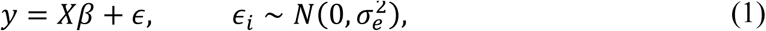

where *y* is an *n*-vector of phenotypes; *X* is an *n* by *m* matrix of genotypes; *β* is an *m*-vector of effect sizes; and *ϵ* is an *n*-vector of residual errors; each element *ϵ_i_* is independent and identically distributed from a normal distribution with variance 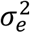. We center the phenotype *y* and standardize each column of the genotype matrix *X* to have zero mean and unit variance, allowing us to ignore the intercept in the model.

Because *p ≫ n,* we have to make further modeling assumptions on the SNP effect sizes *β* to make the model identifiable. We do so by incorporating SNP annotation information and making the effect size β_j_ of *j*-th SNP depending on its annotations. Specifically, we assume that all SNPs are characterized by a same set of *c* annotations. For the j-th SNP, we denote *A_j_ =* (1*,C_j_*_1_*,C_j_*_2_*,…,C_jc_*)^*T*^ as a (*c*+*1*)-vector of realized annotation values including a value of one that corresponds to the intercept. These annotations can be either discrete or continuous. For example, one annotation could be a binary indicator on whether the SNP resides in exonic regions, while another annotation could be a continuous value of the CADD score [18] of the SNP. Because our model includes an intercept (more details in the next paragraph), we require that any linear combination of these annotations does not sum to a vector of one’s across SNPs in order to avoid identifiability issues – a requirement holds for standard linear regression models. For example, we cannot include two non-overlapping annotations that form a partition of the genome (i.e. *C_j_*_1_ *=* 1 when *C_j_*_2_ *=* 0, and *C_j_*_1_ *=* 0 when *C_j_*_2_ *=* 1). Our coding scheme is conventionally referred to as the reference coding scheme. To simplify presentation, we assemble the annotation vectors across all SNPs into an *m* by (*c+1*) annotation matrix *A*, where each row contains the annotation vector for the corresponding SNP.

We assume that the annotations for a given SNP influence its effect size. In particular, we assume that each effect size *β_j_* is independent and follows a normal distribution with mean zero and a variance 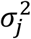 that is SNP specific. The SNP-specific variance assumption generalizes the standard LMM assumption where every SNP is assumed to share a common variance [37,52]. We further impose an extra layer of hierarchy by assuming that the SNP specific variance is a function of the annotation vector, or

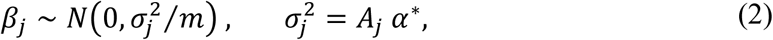

where 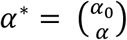 is a (*c+1*)-vector of coefficients that include an intercept *α*_0_ and a *c*-vector of annotation coefficients *α.* Each annotation coefficient is large when the corresponding annotation is predictive of the SNP effect size. Therefore, the annotation coefficients can be used to evaluate the importance of annotations. Above, we center the 2^nd^ to the (c+1)-th columns of the annotation matrix *A* to have mean zero across SNPs. After centering, the ratio 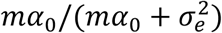 has the natural interpretation of SNP heritability, which is defined as 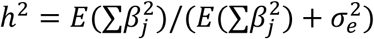, roughly following [16], where *E* represents prior expectation. The intercept *α*_0_ effectively determines how large a typical SNP effect size would be, while the other annotation coefficients determine how the SNP effect size variance would vary around the average depending on what annotations the SNP has. Note that the assumed linear relationship between the SNP specific variance and annotations also naturally extends the modeling assumptions made in LDSC [16] and MQS [48], both of which examine one annotation at a time in the presence of multiple binary annotations, though LDSC has also been recently extended to examine one annotation at a time in the presence of continuous annotations [31]. In addition, our polygenic modeling assumption complements alternative approaches in using sparse models for integrating functional annotations [11,12,40,53].

For inference on the annotation coefficients (*α*), we follow the main idea of LDSC and MQS in using the marginal *χ*^2^ statistics. Using marginal statistics allows our method to be applied to data where only summary statistics are available. Unlike the detailed algorithms of LDSC or MQS that were initially designed to examine one annotation at a time in the presence of multiple binary annotations, however, we applied the generalized estimating equation (GEE) [46,54] inference method that allows for the joint inference of multiple binary and continuous annotations (details in Supplementary Text). GEE is widely used for modeling correlated data and is particularly suitable here to account for the correlation among the marginal *χ*^2^ statistics due to linkage disequilibrium. In the case of binary annotations, the results of our GEE on each annotation by using a diagonal matrix as the working covariance matrix can reduce to that of LDSC and MQS, while the results of our GEE by using an LD based general working covariance matrix can reduce to that of polyGEE [47]. Importantly, just like other summary statistics based methods, GEE inference can be carried out using summary statistics that include marginal *χ*^2^ statistics and the *m* by *m* SNP correlation matrix. The SNP correlation matrix can be obtained from a reference panel, by using, for example, the genotypes from the 1,000 Genomes Project [55]. To facilitate both computation and memory storage, we further approximate the SNP correlation matrix by a block diagonal matrix (details in Supplementary Text), allowing us to capture the main block-wise linkage disequilibrium pattern commonly observed in the human genome [40,56–58]. Finally, with GEE, we obtain both point estimates 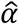 and their variance 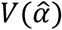 for all annotation coefficients in a closed form. We can then compute the multivariate Wald statistic 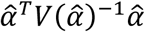 which can be used as a measurement of trait-tissue relevance.

Previous approaches to identify trait-relevant tissues examines one (univariate) Wald statistics at a time, and uses an asymptotic normal test to obtain a p-value to identify significant trait-tissue pairs. Because one annotation in one tissue is often highly correlated with the same annotation in other tissues as well as other annotations in other tissues, the p-values for even the trait-irrelevant tissues are often significant due to the annotation correlation across annotations and tissues. Indeed, as previous studies have shown, even in simple simulations, trait-irrelevant tissues can be falsely identified as trait-relevant in 20% of the simulation replicates [16]. As a consequence, previous studies have to use a set of baseline annotations as covariates to reduce the cross-tissue correlation among annotations, thus reducing false positives. However, it is often unclear how many and what types of baseline variables one should include for a given data set: using a small number of baseline covariates may not control for false positives well, while using a large number of covariates may reduce the power to detect the true trait-relevant tissues. Indeed, the use of baseline variables seems to be highly dependent on data sets (with varying sample sizes and SNP numbers), and needs adjustment in different data sets to achieve sensible results [16].

Here, we present an alternative strategy for identifying trait-relevant tissues. Specifically, for each trait in turn, we model the multivariate Wald statistics across tissues with a mixture of two non-central chi-squared distributions to classify tissues into two groups. The two non-central chi-squared distributions have the same degrees of freedom that equals to the number of annotations fitted in GEE (i.e. *c*), but different noncentrality parameters. The chi-squared distribution with the small noncentrality parameter represents the empirical null distribution that contains tissues irrelevant to the trait. The small, nonzero, noncentrality parameter characterizes the fact that these irrelevant tissues tend to have Wald statistics larger than what would be expected under the theoretical null distribution (i.e. central chi-squared) simply due to annotation correlation across tissues. In contrast, the chi-squared distribution with the large non-central parameter represents the alternative model that contains tissues relevant to the trait. The large noncentrality parameter characterizes the fact that these relevant tissues tend to have Wald statistics larger than those from the irrelevant tissues. By classifying tissues into two groups, we can identify tissues with strong trait-relevance without the need to explicitly model the empirical null distribution using a data generative model. Therefore, our strategy effectively formulates the task of identifying trait-relevant tissues as a classification problem instead of a testing problem. By modeling the empirical null distribution directly, we can reduce false discoveries and potentially gain power at a given false discovery rate (FDR). We also note that this classification strategy follows closely recent applications of mixture models to estimate the empirical null distribution in other genomics settings [59,60]. Technically, we use the expectation-maximization (EM) algorithm to fit the mixture model and infer the two noncentrality parameters as well as the proportion of alternatives from data at hand (details in the Supplementary Text). For each tissue in turn, we then obtain the inferred posterior probability (PP) of it being in the alternative model as its evidence for trait-relevance. We use these inferred posterior probabilities (ranging between 0 and 1) for all following analyses. Note that while our linear mixed model itself does not explicitly model the correlation structure among annotations across tissues by incorporating all annotations from all tissues into a single model, our mixture model and classification strategy can implicitly account for the annotation correlation across tissues.

Finally, we ask the question of how to make use of the inferred trait-relevant tissues to enable more powerful future association studies. We note that our model defined in equations (1) and (2) is closely related to the sequence kernel association test (SKAT) model [49–51] for SNP set test. In particular, the SNP specific variance 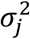 in our model can be naturally interpreted as the SNP specific weight in the SKAT [49–51] framework. Because of this close relationship between our model and SKAT [49–51], we propose to use the estimated SNP specific variance 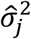 in the top trait-relevant tissue from our model as SNP weights to perform SKAT [49–51] analysis in new association studies. Intuitively, if a SNP tends to have a large effect size, then weighting it high in the subsequent SNP set analysis can help achieve greater association mapping power. We examine this intuition with both simulations and real data applications. Note that our weighted SKAT [49–51] approach is related to a recent method, FST, which also extends SKAT to accommodate multiple functional annotations [32]. However, our method borrows information across all SNPs to infer trait-relevant tissues and estimate annotation coefficients, and further relies on the estimated annotation coefficients in the trait-relevant tissues to construct SNP specific weights for SKAT analysis. In contrast, FST [32] examines one gene at a time, calculates a test statistic for each annotation in turn, and effectively chooses the minimal test statistics among all these annotations as the final statistics for testing. In addition, while our method is polygenic in nature, the idea of using SNP specific weights to construct test statistics is also related to a recent study that uses functional annotations to design SNP specific weights in sparse regression models to improve disease risk prediction performance [61].

### Tissue-Specific SNP Annotations

We used tissue-specific SNP annotations from the ENCODE [23] and the Roadmap [22] projects in the present study. Specifically, we downloaded the broadPeak files from the Roadmap Epigenomics web portal (http://egg2.wustl.edu/roadmap/web_portal/). The broadPeak files contain peak regions for four histone marks (H3K27me3, H3K36me3, H3K4me1, H3K4me3) from 16 cell lines in the ENCODE project and 111 tissues from the Roadmap project (release 9). Among the 127 tissue/cell types, we excluded ESC, IPSC, and ES-derived cell lines to focus on the remaining 105 tissue/cell types (Table S1). Following previous studies [16,22,27], we further classified 105 tissues into 10 tissue groups based on anatomy (BloodImmune, Adipose, AdrenalPancreas, BoneConnective, Cardiovascular, CNS, Gastrointestinal, Liver, Muscle, Other; Table S1). For each tissue and each histone mark in turn, we created a binary histone mark annotation indicating whether the SNP resides inside the peak regions of the histone mark. The average proportions of SNPs residing in each of the four mark labeled regions across the 105 tissues are 25.75% for H3K27me3, 18.51% for H3K36me3, 17.98% for H3K4me1, 10.69% for H3K4me3 (Table S1). In addition to the binary annotations, for each tissue group and each histone mark in turn, we averaged the binary annotation indicator across all tissue types within the tissue group and used the average value as a new, continuous, tissue group level histone mark annotation. Therefore, we obtained both tissue-specific binary histone mark annotations and tissue-group-specific continuous histone mark annotations.

Besides the above histone mark annotations, we also obtained SNP annotations based on 15 chromatin states (TssA, TssAFlnk, TxFlnk, Tx, TxWk, EnhG, Enh, ZNFRpts, Het, TssBiv, BivFlnk, EnhBiv, ReprPC, ReprPCWk and Quies) inferred from ChromHMM [62] in the 105 tissues. In particular, we downloaded the posterior probabilities of each of the 15 states for each genomic location in different tissues from the Roadmap Epigenomics web portal. For each tissue group and each posterior probabilities in turn, we then averaged the posterior probabilities across all tissue types within the tissue group and used the average value tissue as tissue group specific continuous ChromHMM annotation.

### Simulations

We performed two sets of simulations to illustrate the benefits of our method in terms of (1) using multiple SNP annotations and (2) enabling more powerful SNP set tests. For all simulations, we used real genotypes from the Genetic Epidemiology Research Study on Adult Health and Aging (GERA; dbGaP accession number phs000674.v2.p2) [63,64]. The original genotype data of the GERA study consists of 675,367 SNPs on 62,313 individuals. We removed SNPs with a missingness percentage above 0.05, a minor allele frequency (MAF) below 0.05, and a Hardy-Weinberg equilibrium test p-value below 10^−4^. We then randomly selected 10,000 individuals with European ancestry, and obtained the first 27,640 (or 10,000) SNPs on chromosome one to perform the first (or the second) set of simulations.

For the first set of simulations, we obtained two histone marks (H3K4me1 and H3K4me3) from ten different tissue groups from the ENCODE and Roadmap projects, and used them as SNP annotations (details in the previous subsection). Among the ten tissue groups, we randomly selected one as the trait-relevant tissue group in each simulation replicate. We designated all SNPs to be causal, and simulated the causal SNP effects independently from a normal distribution with a SNP-specific variance determined by annotations in the trait-relevant tissue. In particular, we set the variance intercept (i.e. α_0_) to be 0.1 and we varied each of the two annotation coefficients (i.e. α_1_, *α_2_*) from −0.1 to 0.5 (−0.1/0/0.05/0.1/0.25/0.5) to cover a range of possible values estimated from real data (details in Real Data Applications). We performed 1,000 simulation replicates for each combination of the two annotation coefficients (α_1_, *α*_2_). Note that the median estimates of the two annotations across 43 GWAS traits (details in Real Data Applications) is close to (*α*_1_, *α*_2_)*_m_ =* (0.1, 0.05). We simulated the residual errors from a normal distribution with variance 0.9, so that the resulting trait has a SNP heritability of 0.1, which corresponds to the median SNP heritability estimate across 43 traits in the real data analysis. We then summed all genetic effects and the residual errors together to form the simulated phenotypes. With genotypes and simulated phenotypes, we fitted a linear regression model for one SNP at a time and computed marginal χ^2^ statistics. We further paired these marginal statistics with a SNP correlation matrix estimated using 503 individuals of European ancestry from the 1,000 Genomes Project [55]. We then examined the ten candidate tissues in turn using either two annotations together or one annotation at a time. For additional comparisons at the median setting (α_1_, *α*_2_)*_m_ =* (0.1, 0.05), we also included LDSC [16], which in default includes 75 baseline annotations as covariates. We used this first set of simulations for two purposes. In the Supplementary Text, we used simulations to illustrate the benefits of using mixture models to post-process the Wald statistics in order to address correlations among annotations and reduce false positives (Figure S1). In the main text, we used simulations to illustrate the benefits of modeling multiple annotations jointly.

For the second set of simulations, we used 10,000 SNPs and divided them into 100 blocks with 100 SNPs inside each block. For the null simulations, we set the effect sizes of all SNPs to be zero and performed 50,000 simulation replicates. For the alternative simulations, we randomly selected 10 non-adjacent blocks as causal blocks and we randomly selected 20% SNPs inside these causal blocks to be causal SNPs (i.e. a total of 200 causal SNPs). We then simulated ten tissue-specific annotation sets, each with two annotations, which are simulated to correlate with SNP causality [30]. Specifically, the annotation values for the non-causal SNPs are sampled from a normal distribution with mean 0 and variance 1. The causal SNPs are randomly divided into three groups: for the first annotation, its annotations values for the first group are sampled from a normal distribution with mean 0 and variance 1 while its annotations values for the second and third groups are sampled from a normal distribution with mean 10 and variance 1; for the second annotation, its annotations values for the second group are sampled from a normal distribution with mean 0 and variance 1 while its annotations values for the first and third groups are sampled from a normal distribution with mean 10 and variance 1. The proportion of the three groups of causal SNPs are set to be either (1/2, 1/2, 0), (1/3, 1/3, 1/3) or (0, 0, 1). Because two annotations share similar annotation values in the third group of causal SNPs, the proportion of the third group determines the correlation between the two annotations for causal SNPs within the annotation set. Therefore, the selected proportion of the third group SNPs being 0, 33% and 100% correspond to low, median and high correlation between the two annotations in causal SNPs, respectively. Once we had the annotations, we simulated the effect sizes for causal SNPs independently from a normal distribution with a SNP-specific variance determined by the designated annotation set. Specifically, we set α_0_ = 0.5 and chose either (α_1_, α_2_) = (0.4, 0.4) (in the case of two informative annotations) or (α_1_, α_2_) = (0.4, 0) (in the case of one informative annotation). These parameters were selected to ensure that the 10 causal blocks explain a large proportion of variance in phenotypes (per-block PVE > 0.01; Figure S2) so that we will have reasonable power to detect them. Certainly, power is a continuous function of per-block PVE and is non-zero even for small values of per-block PVE. We simulated the residual errors from a normal distribution with variance 0.5. We summed all genetic effects and the residual errors together to form the simulated phenotypes. We then randomly divided the 10,000 individuals into two sets: a training set with 7,000 individuals and a test set of 3,000 individuals. In the training set, we followed the same procedure described in the previous paragraph to obtain marginal χ^2^ statistics in the data and SNP correlation matrix from a reference panel to fit our model. We applied the parameter estimates from the best trait-relevant tissue determined in the training set to compute the SNP specific variance 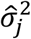 as SNP weights. For the computed variance, we subtracted from them the minimal variance across SNPs and added a small constant (10^−15^) to ensure that all weights are positive. We then multiplied the resulting SNP weights with the posterior probability (PP) of the best trait-relevant tissue and further added a value of 1-PP to all SNPs, thus effectively obtaining a set of averaged SNP weights by using both the constructed SNP weights in the identified trait-relevant tissue and the equal SNP weights. Averaging the constructed weights and the equal weights using PP ensures the robustness of our method and guards against the special case where none of the tissues are trait-relevant: in this case, the resulting SNP weights would equal to the equal weights due to a small PP value and would thus still be effective in the subsequent SNP set analysis. We finally applied the SNP weights constructed in the training data to the test data to perform SNP set analysis. We performed 1,000 simulation replicates for each alternative simulation setting. We divided these replicates into 10 sets, each with 100 replicates, and computed the power to detect the causal blocks in each set. We report the mean and variance of these power values across 10 sets.

### GWAS Summary Statistics

We obtained summary statistics in the form of marginal z-scores for 43 traits from 28 GWAS studies. Details are provided in Table S2. These studies collect a wide range of complex traits and diseases that can be classified into six phenotype categories [28,65]: anthropometric traits (e.g. height and BMI), hematological traits (e.g. MCHC and RBC), autoimmune diseases (e.g. CD and IBD), neurological diseases (e.g. Alzheimer’s disease and Schizophrenia), metabolic traits (e.g. FG and HDL), and social traits (e.g. ever smoked and college completion). We removed SNPs within the major histocompatibility complex (MHC) region (Chr6: 25Mb-34Mb) following [16]. We then intersected the SNPs from all the studies and retained a common set of 622,026 SNPs for analysis. We paired the marginal z-scores from these studies with the SNP correlation matrix estimated using 503 individuals of European ancestry from the 1,000 Genomes Project [55] for inference. Finally, after the analysis, we computed correlation among traits in terms of their tissue relevance and used the Bayesian information criterion (BIC) implemented in the clustering package mclust [66] in R with the standard option EEI to classify traits. Clustering with BIC automatically inferred a total of five main trait clusters.

### Investigate Trait-Tissue Relevance via PubMed Search

We rely on previous literature to partially validate the inferred trait-relevant tissue results in real data. We reasoned that, if a tissue is indeed relevant to a given trait, then there would be extensive prior biomedical researches carried out on the tissue for the trait. Therefore, the number of previous publications on a trait-tissue pair can serve as a useful indicator on the potential relevance and importance of the tissue for the trait. To estimate the number of previous publications on trait-tissue pairs, we conducted a literature search on PubMed (https://www.ncbi.nlm.nih.gov/pubmed/). Specifically, for each trait-tissue pair, we used the names of trait and tissue as input and counted the number of publications that contain the input values either in the abstract or in the title. For traits, we used trait names directly. For tissues, we excluded the “Other” tissue group and focused on the nine remaining tissue groups. For these remaining tissue groups, we used the following key words in addition to the tissue group name in the PubMed search: (1) CNS: brain, central nervous system, neuron, glia and CNS; (2) BloodImmune: blood, T-cell, B-cell, thymus and immune system; (3) Adipose: adipose. (4) AdrenalPancreas: adrenal, pancreas; (5) BoneConnective: bone, fibroblast and connective tissue; (6) Cardiovascular: heart, cardiovascular; (7) Gastrointestinal: gastrointestinal, esophagus, stomach, intestine and rectum; (8) Liver: liver; (9) Muscle: muscle. For example, for the schizophrenia-CNS trait-tissue pair, we conducted the search by using “schizophrenia [Title/Abstract] AND (CNS [Title/Abstract] OR brain [Title/Abstract] OR central nervous system [Title/Abstract] OR neuron [Title/Abstract] OR glia [Title/Abstract])”, which yielded 17,720 hits. The number of publications on each trait-tissue pair from the PubMed search is listed in Table S3 (the search was carried out on June 23, 2017). For each trait in turn, we further normalized the data by dividing the number of publications for a tissue by the total number of publications across all tissues for the trait. We used the resulting proportion for the final analysis.

### SNP Set Test in WTCCC

Because of the close relationship between our method and the sequence kernel association test (SKAT [49–51]) (details in Method Overview), we propose to use the estimated SNP specific variance in the top trait-relevant tissue from our method as SNP weights in SKAT [49–51] to perform SNP set test in new association studies. To examine the utility of this association mapping strategy in real data, we estimated annotation coefficients in consortium studies, applied them to construct SNP weights (details in the above Simulations subsection), with which we performed SKAT [49–51] for the corresponding traits in the Wellcome Trust Case Control Consortium (WTCCC) study [67]. The WTCCC data consists of about 14,000 cases from seven common diseases and 2,938 shared controls. The cases include 1,963 individuals with type 1 diabetes (T1D), 1,748 individuals with Crohn’s disease (CD), 1,860 individuals with rheumatoid arthritis (RA), 1,868 individuals with bipolar disorder (BD), 1,924 individuals with type 2 diabetes (T2D), 1,926 individuals with coronary artery disease (CAD), and 1,952 individuals with hypertension (HT). We excluded HT and considered the remaining six diseases for which we had summary statistics in other larger consortium studies. We obtained quality controlled genotypes from WTCCC [67] and imputed missing genotypes using BIMBAM [68] to obtain a total of 458,868 SNPs that are shared across all individuals. The genotypes were further imputed by SHAPEIT [69,70] and IMPUTE2 [71] with the 1,000 Genomes Project [55] as a reference. We removed SNPs with a Hardy-Weinberg equilibrium p-value < 10^−4^ or a minor allele frequency < 0.05, and intersected SNPs from WTCCC with the consortium data to obtain a final set of 335,170 overlapped SNPs. Meanwhile, we obtained genome locations for a set of 19,189 protein coding genes from GENCODE project [72]. We intersected SNPs with these genes and identified gene-harboring SNPs that reside within 10 kb upstream of the transcription start site and 10 kb downstream of the transcription end site. To perform gene-set test, we focused on 5,588 genes that have at least 10 SNPs, with an average of 29.6 SNPs inside each gene and a total of 153,813 gene-harboring SNPs. For each gene in turn, we computed SNP-specific variance using annotation coefficient estimates from the best trait-relevant tissue inferred with consortium study summary statistics for the corresponding trait. We used the SNP-specific variance as SNP weights. As in simulations, for these weights, we subtracted from them the minimal weight across SNPs and added a small constant (10^−15^) to ensure that all weights are positive. We then multiplied the resulting SNP weights with the posterior probability (PP) of the best trait-relevant tissue and further added a value of 1-PP to all SNPs as the final SNP weights to perform SKAT [49–51] analysis. The SNP-weights for the 43 traits can be downloaded from http://www.xzlab.org/.

## Results

### Simulations: Multiple Annotations versus Single Annotation

Our first set of simulations are used to illustrate the benefits of using multiple annotations to identify trait-relevant tissues. Details of simulations are provided in Materials and Methods. Briefly, we obtained 27,640 SNPs from 10,000 randomly selected individuals in the GERA study [63,64] and simulated phenotypes. We considered two histone annotations (H3K4me1, H3K4me3) from ten tissue groups (Table S1), among which we randomly designated one as the trait-relevant tissue. We then simulated SNP effect sizes under a polygenic model based on the two annotations in the trait-relevant tissue. We added genetic effects with residual errors to form simulated phenotypes. We obtained summary statistics from the data and considered three different approaches to identify trait-relevant tissues:

1. SMART. We analyzed two annotations jointly and computed a single multivariate Wald statistic for each tissue using our procedure. We then applied an EM algorithm and a mixture model on the multivariate Wald statistics to further classify tissues into two groups. We used the posterior probability of a tissue being trait-relevant to measure trait-tissue relevance.
2. Uni. We analyzed one annotation at a time and computed two univariate Wald statistics for each tissue using our procedure. We then applied an EM algorithm to classify these Wald statistics into two groups. For each tissue and each annotation, we obtained the posterior probability of being a trait-relevant tissue to measure trait-tissue relevance.
3. UniMax. We analyzed one annotation at a time. For each tissue, we computed two univariate Wald statistic using our procedure and selected among them the larger statistic as a measurement of trait-tissue relevance. We then applied an EM algorithm to classify these Wald statistics into two groups. For each tissue, we obtained the posterior probability of its being a trait-relevant tissue to measure trait-tissue relevance.

Above, we have included two versions of univariate tests: Uni and UniMax. While the Uni approach is widely applied in previous studies [16,27,28,48], the UniMax approach can be statistically more appropriate than Uni for summarizing tissue-level evidence. We considered a range of realistic annotation coefficient combinations (i.e. (*α*_1_, *α*_2_)). For each combination, we performed 1,000 simulation replicates. For each method, we computed the power of various methods in detecting the trait-relevant tissue at a false discovery rate (FDR) of 0.1 (Figure 1A). In the majority of settings, analyzing multiple annotations jointly also improves power compared with analyzing one annotation at a time. For example, based on power at 10% FDR, SMART is ranked as the best method in 15 out of 25 simulation settings where both annotations have non-zero effects, while UniMax is the best in 10 settings (Figure 1A). While the performance of SMART is often followed by UniMax, the power improvement of SMART compared with UniMax can be large (median improvement = 9.2%). Certainly, in the special cases where one annotation coefficient is exactly zero or close to zero, then SMART is often outperformed by UniMax, presumably due to its smaller degree of freedom there. For example, among the 11 settings where at least one annotation has zero effects (grey area, Figure 1A), SMART is ranked as the best method only 4 times, while UniMax is ranked as the best method 7 times. Finally, to further explore the characteristics of annotations that can influence the power of SMART in identifying trait-relevant tissues, we simulated annotations that have various genome-occupancy characteristics and that have various annotation effect sizes and signs (Supplementary Text). We show that the power of SMART increases with increasing annotation coefficients, is not sensitive to the signs of annotations, and is relatively stable with respect to the genome-occupancy of the annotations as we have standardized the annotations to have mean zero and standard deviation one across the genome (Figure S3).

**Figure 1.**
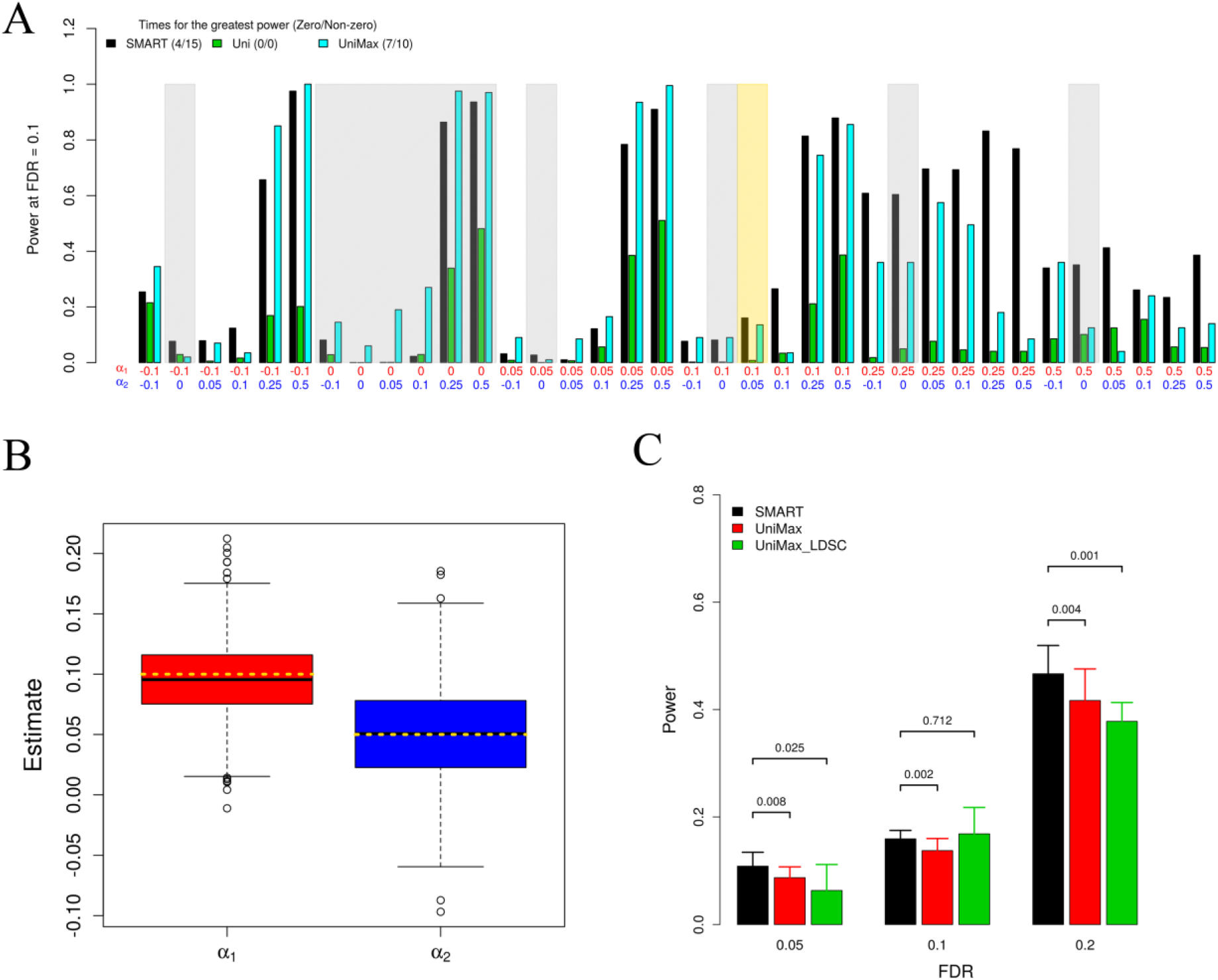
Simulation results for comparing using multiple annotations versus a single annotation. (A) Power to detect trait-relevant tissues by different approaches in various settings at a fixed FDR of 0.1. x-axis shows the values of the two annotation coefficients used in the simulations. Settings where at least one annotation coefficient is zero are shaded in grey. The setting where the annotation coefficients equal to the median estimates from real data (i.e. ***α*** = (**0.1, 0.05**)) is shaded in gold. The first number for each method in the figure legend represents the number of times each method is ranked as the best in 25 simulation settings where none of the annotations have zero coefficients; while the second number represents the number of times each method is ranked as the best in 11 simulation settings where at least one annotation has a zero coefficient. (B) Annotation coefficient estimates by SMART are centered around the truth (horizontal dotted gold lines). (C) Mean power (y-axis) to detect trait-relevant tissues by different approaches at different FDR values (x-axis). Error bar shows the standard deviation computed across 10 simulation groups, each of which contains 1,000 simulation replicates (i.e. a total of 10,000 simulations). *p*-values from the paired t-test are used to compare methods at different FDR cutoffs. Note that the error bar is large due to the small number of simulation replicates within each simulation group. For (B) and (C), simulations were done at ***α*** = (**0. 1, 0. 05**). FDR, false discovery rate.

We examine in detail a simulation setting where (*α*_1_, *α*_2_) are chosen to be close to the median estimates (0.1, 0.05) from the real data sets (i.e. gold shade in Figure 1A and Figure S1). Note that even though these parameters are chosen based on real data, we have much less SNPs or samples in the simulations than in real data and are thus underpowered in simulations. In any case, we first obtained annotation coefficient estimates 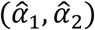 across simulation replicates in this setting. We found that the estimates are centered around the truth as one would expect (Figure 1B), suggesting accurate parameter estimation by our approach. Next, in addition to the six approaches listed above, we also included a UniMax_LDSC approach into comparison. In the UniMax_LDSC approach, we applied LDSC to analyze one trait at a time and used the maximum Wald statistics among the two to measure trait-tissue relevance. Different from the UniMax_Wald, however, UniMax_LDSC used a set of 75 baseline annotations to address correlation among annotations. As a result, UniMax_LDSC performs similarly as UniMax in terms of power to detect trait relevant tissues at different FDR thresholds (Figure 1C), suggesting that using mixture modeling is competitive compared to using covariates to control for annotation correlation across tissues. Because both UniMax_LDSC and UniMax use only one annotation, they are often less powerful compared to SMART that models two annotations together (Figure 1C).

### Simulations: Construct Powerful SNP Set Test

Our second set of simulations is intended to illustrate the benefits of our method in using inferred trait-relevant tissue to enable more powerful SNP set tests. Here, we ask the question of how to make use of the inferred trait-relevant tissues to enable more powerful future association studies. As explained in the Method Overview section, our model is closely related to the sequence kernel association test (SKAT) [49–51] for SNP set test. In particular, the SNP specific variance in our model can be naturally interpreted as the SNP specific weight in the SKAT [49–51] framework. Because of this close relationship between our model and SKAT [49–51], we propose to use the estimated SNP specific variance in the top trait-relevant tissue from our model as SNP weights to perform analysis in new association studies. Intuitively, if a SNP tends to have a large effect size, then weighting it high in the subsequent SNP set analysis can help achieve greater association mapping power. To explore the possibility of using the inferred SNP-specific variance estimate 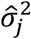 as *a priori* weight to construct SNP set test in the SKAT framework [49–51], we obtained 10,000 SNPs from the same set of 10,000 individuals in the GERA study [63,64] and simulated phenotypes (Materials and Methods). To do so, we divided these SNPs evenly into 100 blocks and randomly selected 10 blocks to be causal blocks. In each casual block, we further selected 20 SNPs to be causal SNPs. We then simulated ten tissue-specific annotation sets with two annotations within each set and designated one set as the trait-relevant tissue. We simulated causal SNP effect sizes based on the two annotations from the trait-relevant tissue and added residual errors to form the simulated phenotypes. Afterwards, we divided individuals randomly into two sets: a training set of 7,000 individuals and a test set of 3,000 individuals. We applied SMART_EM and UniMax_EM in the training set to identify trait-relevant tissues and to estimate annotation coefficients. We then applied the following weighting options to perform SKAT [49–51] analysis in the test set:

1. EqualWeight, where we weight all SNPs equally.
2. TissueWeight_Oracle, where we use the true coefficients from the correct trait-relevant tissue to construct SNP weights. This represents an up limit of power we can possibly achieve.
3. TissueWeight_SMART, where we fitted SMART_EM in the training data and applied the coefficient estimates for the two annotations in the inferred trait-relevant tissue to construct SNP weights.
4. TissueWeight_UniMax, where we fitted UniMax_EM in the training data and applied the coefficient estimate for the annotation with the larger Wald statistics in the inferred trait-relevant tissue to construct SNP weights.

We first simulated 50,000 replicates under the null where there is no causal SNP so that both α_0_ and (α_1_, α_2_) are 0. We used the null simulations to examine the type I error control for various methods and we found that all these methods are well behaved (Figure 2A). Next, we simulated 1,000 replicates under the alternative where we have non-zero α_0_ and (α_1_, α_2_). We divided SNPs into 100 blocks, among which 10 are causal. We compared different methods in terms of their power to identify the causal blocks. In the simulations, we generated two annotations whose values in the trait-relevant tissue are predictive of SNP causality. The annotation values for the two annotations are almost identical in a certain proportion of causal SNPs (chosen to be 0%, 33%, or 100%) so that the two annotations can contain complementary information (in the case of 0%) or overlapping information (in the case of 100%). Intuitively, information overlapping in the two annotations would reduce the relative power gain of using multiple annotations versus using a single annotation in constructing SNP set tests. As one would expect, in all settings, constructing SNP weights using the true coefficients from the correct trait-relevant tissue (i.e. TissueWeight_Oracle; red bars in Figure 2B and C) achieves the greatest power compared with the other methods. For example, compared with using equal weights, using the oracle weights improves power by 14.1% on average (median = 13.7%) across all settings. Importantly, constructing SNP weights using the estimated coefficients from the inferred trait-relevant tissue weight (i.e. TissueWeight_SMART; green bars in Figure 2 B and C) can often achieve almost identical power as TissueWeight_Oracle. Comparing between TissueWeight_SMART and TissueWeight_UniMax, when both annotations have non-zero coefficients, we found that using multiple annotations often leads to greater power gain than using a single annotation. However, as one would expect, when the two annotations contain overlapping information (e.g. in the case of 100%), then using one annotation yields similar power as using two annotations (green vs blue in Figure 2B). In the special case where only one annotation has a non-zero coefficient, then using multiple annotations also has similar power compared with using a single annotation, even when the two annotations contain complementary information (green vs blue in Figure 2C).

**Figure 2.**
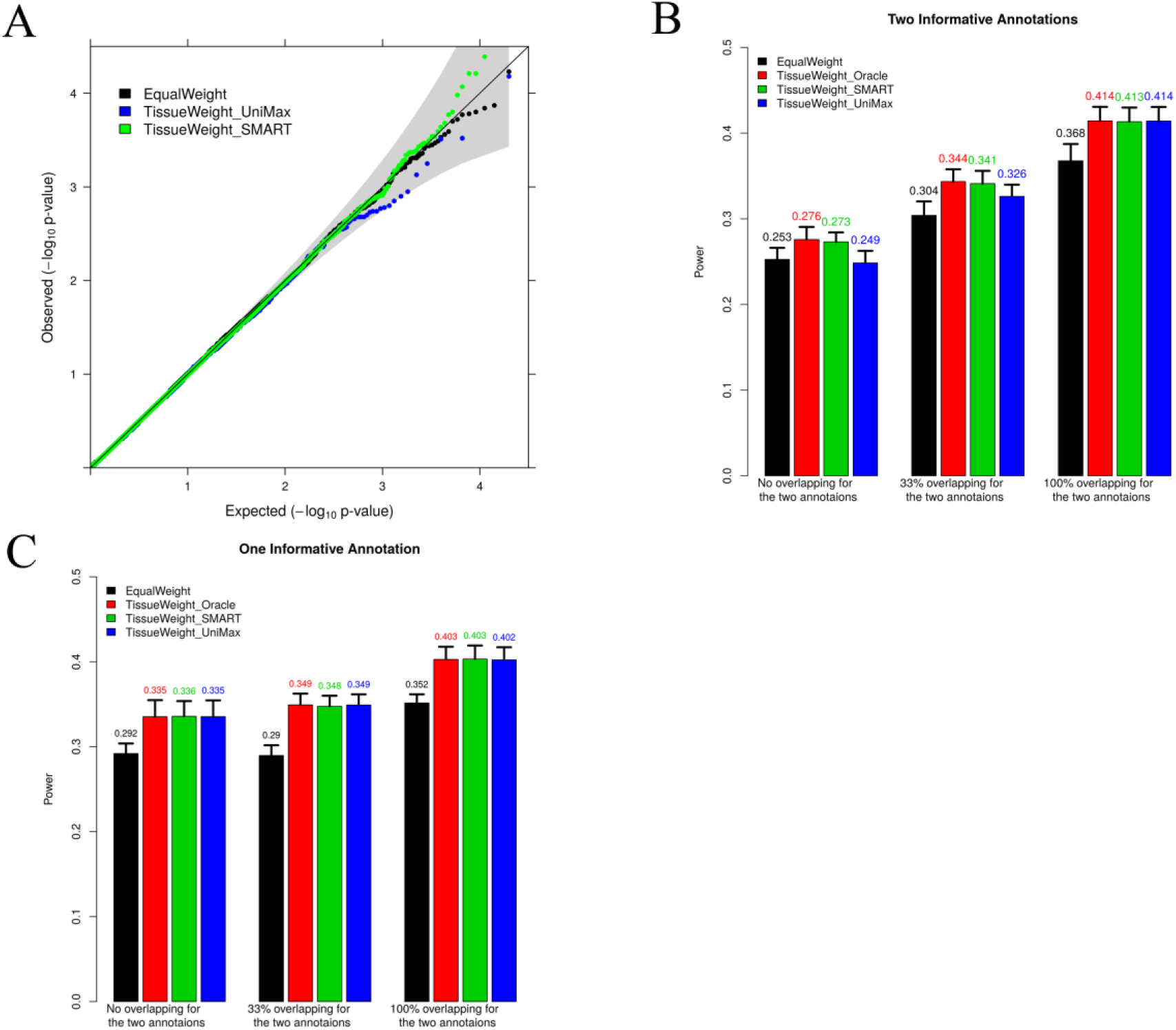
Simulation results for using different weights to construct SNP set tests. (A) QQ plot of −log10 p values from SNP set tests using different SNP weights under the null simulations. Tests using different weights all control type I error well. (B) Power to detect causal blocks by SNP set tests using different SNP weights in the simulation setting where **α** = (**0.4, 0.4**). (C) Power to detect causal blocks by SNP set tests using different SNP weights in the simulation setting where *α* = (**0.4,0**). For both (B) and (C), Power are evaluated at a genome-wide significance threshold of 1×10^−4^. Standard errors are computed across 1,000 simulation replicates. The x-axis shows the proportion of causal SNPs that have identical values for the two annotations, which measures correlation between the two annotations.

Next, we explore how various simulation parameters influence the weighted SKAT power (Supplementary Text). Here, we also include TissueWeight_UniMaxLDSC, where we applied LDSC with the UniMax procedure in the training data and used the coefficient estimate for the annotation with the larger Wald statistics in the inferred trait-relevant tissue to construct SNP weights. In this set of simulations, we varied the annotation coefficients and varied the number of causal blocks. The results are presented in Supplementary Figure S4. With simulations, we show that the power of SNP set test primarily depends on the phenotype variance explained (PVE) by each causal block (i.e. per-block PVE) as well as annotation coefficients, and increases with increasing per-block PVE or annotation coefficients. In contrast, power is not influenced by the number of blocks when per-block PVE is fixed to be a constant, though it would decrease with increasing number of blocks when the total PVE is fixed (as per-block PVE is negatively correlated with number of blocks in this case). Importantly, TissueWeight_SMART outperforms TissueWeight_UniMax and TissueWeight_UniMaxLDSC in most scenarios and outperforms EqualWeight in all scenarios.

## Real Data Applications

### Data processing and analysis overview

We applied our method to analyze 43 traits from 29 GWAS studies (Table S2). For these analyses, we obtained tissue-specific SNP annotations in 105 tissues from 10 tissue groups in the ENCODE [23] and Roadmap [22] projects and processed these annotations into three sets (details in Materials and Methods; Table S1): (1) four binary annotations (H3K27me3, H3K36me3, H3K4me1, and H3K4me3) in 105 tissues constructed based on histone mark occupancy; (2) four continuous annotations in 10 tissue groups (BloodImmune, Adipose, AdrenalPancreas, BoneConnective, Cardiovascular, CNS, Gastrointestinal, Liver, Muscle, and Other) based on averaging binary annotations across tissues within each tissue group; and (3) posterior probabilities of the 15 ChromHMM [62] states in the 10 tissue groups. For each trait in turn, we paired the three sets of annotations with GWAS summary statistics to identify trait-relevant tissues. Data processing and analysis details are described in the Materials and Methods.

### Trait tissue relevance determined by SMART is largely consistent with PubMed literature and highlights the importance of modeling multiple annotations

Besides applying our method to infer trait relevant tissues, we first rely on the knowledge gained from previous biomedical literature to measure trait-tissue relevance. To do so, we conducted a PubMed literature search and counted the number of publications existed for each trait-tissue pair. We reasoned that, if a tissue is indeed relevant to a given trait, then there would be a fair number of studies performed on the tissue for the given trait. Therefore, the number of publications carried out on a trait-tissue pair would be a good indicator on their relevance. Next, for each trait in turn, we normalized the count data by computing the proportion of previous publications performed on each of the nine tissue groups (i.e. the ten tissue groups minus the “Other” group), resulting in for each trait a vector of nine proportion values that sum to one (details in Materials and Methods). The number of total publications and proportion values for all traits are summarized in Table S4. The PubMed literature search results are generally consistent with what we would expect. For example, for schizophrenia (SCZ), 63.8% of the previous literatures are focused on CNS, with the rest of the literature scattered across other tissues. The proportion of literature carried out on each trait-tissue pair obtained in PubMed thus provides a reasonable *a priori* measure of trait-tissue relevance. We use these measurements to validate some of our analysis results using tissue specific annotations.

We then applied our method to jointly analyze multiple annotations for each of the three sets of tissue-specific annotations described above. We denote the analysis on annotation set (1) as ***HB*** (i.e. histone marks, binary), analysis on annotation set (2) as ***HC*** (i.e. histone marks, continuous), and analysis on annotation set (3) as ***cHMM*** (i.e. ChromHMM annotation). For ***HC*** and ***cHMM***, we obtained the posterior probability values (PPs) from each of the 10 tissue groups for each trait. For ***HB***, we first obtained PPs from each of the 105 tissues for each trait. For each trait in turn, we then averaged these PP values within each tissue group and used the averaged tissue group level PP values for the following analyses – this way we can perform comparisons at the tissue group level across different annotation sets. As a comparison, we also applied our method to each of the three annotation sets and performed univariate analysis that corresponds to the UniMax procedure explained in the simulation section. These univariate analyses include ***HBuMax***, which is a univariate analysis of the annotation set (1); ***HCuMax***, which is a univariate analysis of the annotation set (2); ***cHMMuMax***, which is a univariate analysis of the annotation set (3); and ***HBuMaxLDSC***, which is a univariate analysis on the binary annotation set (1) using LDSC. In these univariate analyses, for each trait in turn, we first selected the annotation with the maximum Wald statistic in every tissue (or tissue group). We then computed the PPs of the selected mark in all tissues (or tissue groups). When necessary, we further averaged PPs (for ***HBuMax***) or Wald statistics (for ***HBuMaxLDSC***) within each tissue group to allow for comparison at the tissue group level. Overall, we obtained 10 tissue group level PP values or Wald statistics for every trait from each of the five different approaches. We list the top trait-relevant tissue groups with the largest PP value or Wald statistics identified by each of the above approaches in Table S3, with the corresponding tissue group PPs listed in the same table. The tissue group PPs from HC are also plotted in Figure S5.

The results from those different approaches are largely consistent with the PubMed search results, though notable deviations exist. For example, PubMed search identified CNS to be the most relevant tissue to five neurological traits (ADD, BIP, SCZ, Autism and Alzheimer). Approaches using annotations also identified CNS as the most trait-relevant tissue for four of the five neurological traits (ADD, BIP, SCZ and Autism). However, for Alzheimer’s disease, using tissue-specific annotations revealed BloodImmune tissue as a trait-relevant tissue, which is consistent with recent discoveries that inflammation and microglia are important for Alzheimer’s disease etiology [73,74]. As another example, PubMed search identified liver to be the most relevant tissue for hematological traits (MCHC, MCH, MCV, MPV, PLT and RBC), presumably because of liver’s important role in producing extrarenal erythropoietin [75]. In contrast, using tissue-specific annotation highlighted BloodImmune tissue as the most relevant tissue for hematological traits. The similarity and difference between SMART and PubMed search results highlight the importance of using different information to infer trait tissue relevance.

We further quantify the similarity between various approaches and PubMed results. To do so, we compare the tissue group level PP values from the annotation integration approaches to the proportion of publications on each tissue group obtained from PubMed search. For each trait and each approach in turn, we computed the correlation between the PP vector for the nine tissue groups and the corresponding proportion values from PubMed search (Figure 3C). We reasoned that, if an approach makes good use of the annotation information, then the trait tissue relevance inferred by this approach would be consistent with the trait tissue relevance measured by PubMed search. Indeed, the Spearman’s rank correlations between different integrative approaches and the PubMed search are reasonable, with an median (average) value of 0.474 (0.420), 0.417 (0.379), 0.283 (0.242), 0.433 (0.397), 0.433 (0.376), 0.417 (0.360) and 0.417 (0.344) for ***HB, HBuMax, HBuMaxLDSC, HC, HCuMax, cHMM*** and *c****HMMuMax***, respectively. The correlation results also suggest that, for the same annotation set, using multiple annotations often yields better performance than using one annotation alone (i.e. ***HB*** vs ***HBuMax*** or ***HBuMaxLDSC***, ***HC*** vs ***HCuMax***, and ***cHMM*** vs ***cHMMuMax***). Finally, comparing different annotation sets, we found that using 15 chromatin states (i.e. ***cHMM*** and ***cHMMuMax***) often result in lower correlation than using the annotations based on histone occupancy, suggesting that post-processing histone occupancy data into chromatin states may lose important trait-tissue relevance information, dovetailing earlier findings [76].

**Figure 3.**
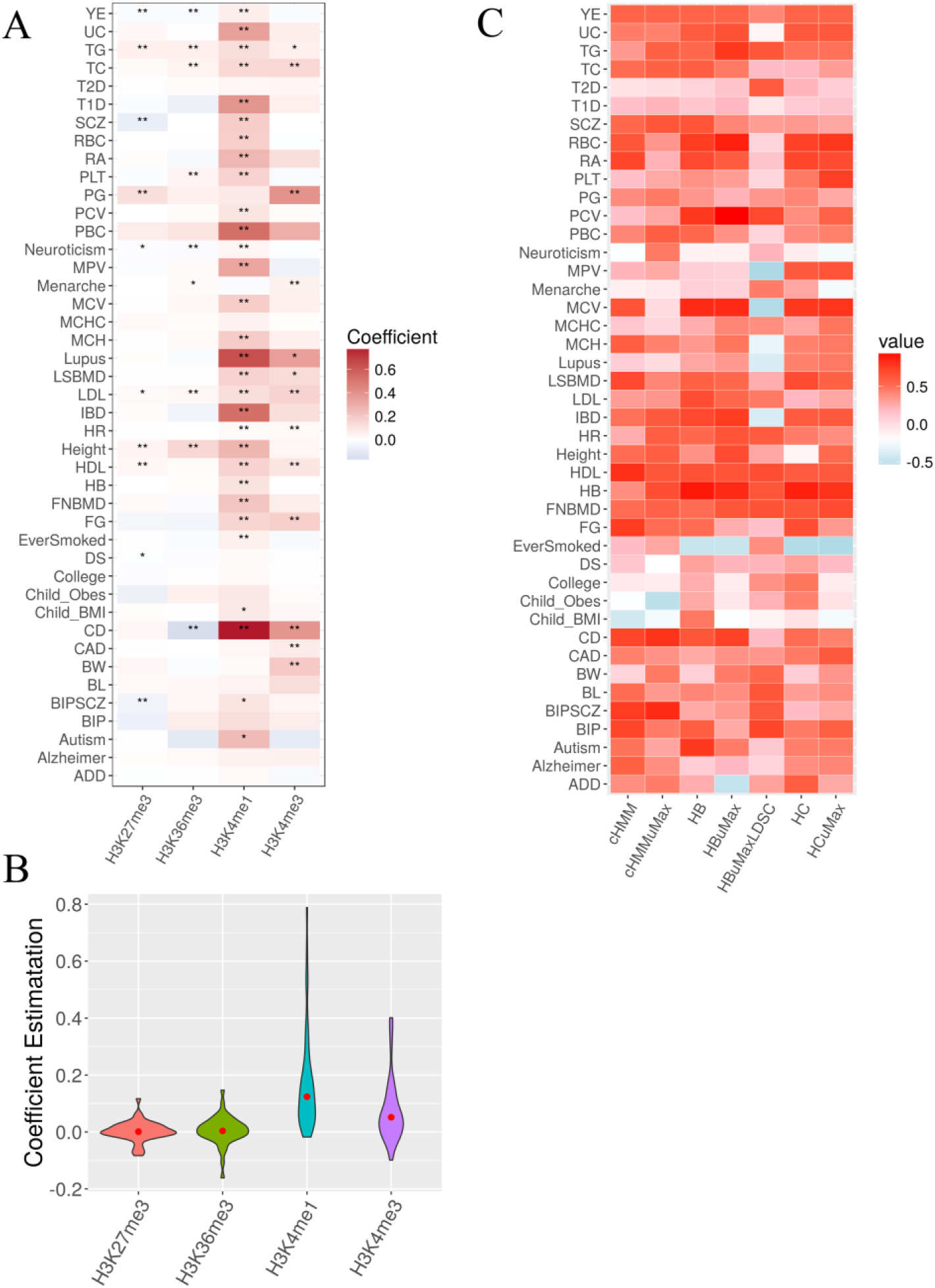
Inferring tissue relevance for 43 GWAS traits using SNP annotations (A) Heatmap displays the annotation coefficient estimates for the four histone marks (y-axis) in the most trait relevant tissue for 43 traits (x-axis). Color captures the sign and magnitude of the estimates, while the number of stars represent the significance of the estimates after Bonferroni correlation for the 43 * 10 *4 = 1,720 hypotheses tested (*: p < 0.05; **: p < 0.01). (B) Boxplot shows the coefficient estimates for the four histone marks in the most trait relevant tissue across 43 traits. (C) Heatmap displays correlation between different annotation integration approaches (y-axis) and the PubMed search approach for 43 GWAS traits (x-axis). For Uni and SMART approaches, correlations are computed between the average posterior probability vectors of the nine tissue groups from different annotation integration approaches and the proportion of existing publications on the same nine tissue groups from PubMed search; For UniMax_LDSC approach, correlations are computed between the average Wald statistic vectors of the nine tissue groups and the proportion of existing publications on the same nine tissue groups from PubMed search. Color captures the sign and magnitude of the estimates.

### Tissue relevance analysis reveals important histone markers in predicting SNP effects and classifies 43 traits into five main clusters

To characterize trait-tissue relevance at the tissue group level, we focused on the results from the ***HC*** approach in more details and examined the annotation coefficients for the four histone marks inferred from the most trait-relevant tissue. We show the estimates and their significance for individual traits in Figure 3A and then grouped coefficients across 43 traits in Figure 3B. Overall, among the four histone marks, two of them have positive coefficient estimates: the coefficient estimates for H3K4me1 are positive for 42 out of 43 traits, while the coefficient estimates for H3K4me3 are positive for 32 traits (Figure 3B). The positive coefficients for the two histone marks are consistent with their role in marking promotors and enhancers, which are enriched in or near association loci identified by multiple GWASs [11,45]. In contrast, the coefficient estimates of H3K27me3 and H3K36me3 are mostly estimated to be close to zero, with approximately random positive or negative signs (positive signs in 22 traits for H3K27me3, and in 25 traits for H3K36me3; Figure 3A). The near-zero estimates of H3K27me3 and H3K36me3 suggests that SNP effect sizes in polycomb repression regions and transcribed regions often do not differ much from the rest of the genome. In terms of the magnitude of the estimated coefficients, two of the four marks (H3K4me3 and H3K4me1) have large effect estimates, as well as high estimation variation, across all examined traits (Figure 3B). The large coefficient estimates for H3K4me3 and H3K4me1 suggest that both promotor regions and enhancer regions are highly predictive of SNP effect sizes and are often the most informative for inferring trait-tissue relevance.

The results with the ***HC*** approach are also consistent with what we see in the simulations. In particular, while our extended linear mixed model itself does not explicitly model the correlation among annotations across tissues, our mixture modeling and classification strategy implicitly accounts for annotation correlation across tissues and allows us to identify multiple trait-relevant tissues for a given trait when they are present. Indeed, examining the tissue group PPs from HC (Figure S5), we found that, among the 43 GWAS traits, more than half of them have one trait-relevant tissue with PP>0.5, while 4 of them have two or more trait-relevant tissues with PPs>0.5. For example, consistent with [16], the CNS tissue group was identified as the trait-relevant tissue for SCZ, BIP, YE and Ever smoked. Consistent with [16], the blood immune tissue group was identified as the trait-relevant tissues for CD, RA and UC. While consistent with [16,27], multiple tissue groups, including bone connective, muscle, cardiovascular and adipose, were identified as relevant for height.

To further characterize trait-tissue relevance at the tissue level, we examined the results from the ***HB*** approach in details. The annotation set (1) contains binary annotations for 105 tissues that belong to 10 tissue groups. We have only focused on examining group-level results from this set of annotations so far. Here, we focus instead on the PP values for the 105 tissues directly; thus we have a 105-vector of PP values for every trait. We relied on the PP values to rank tissues for every trait. The tissue rank list for each trait represents the tissue footprint of each trait: the trait-relevant tissues are ranked high in the list while the trait-irrelevant tissues are ranked low in the list. With the tissue rank list, we assess the similarity between GWAS traits in terms of their tissue relevance by hierarchical clustering (Figure 4). We also computed pair-wise spearman correlation between traits based on the tissue rank list (Figure 5). Overall, applying the Bayesian information criterion (BIC) to the correlation plot using mclust [66] revealed five main trait clusters.

**Figure 4.**
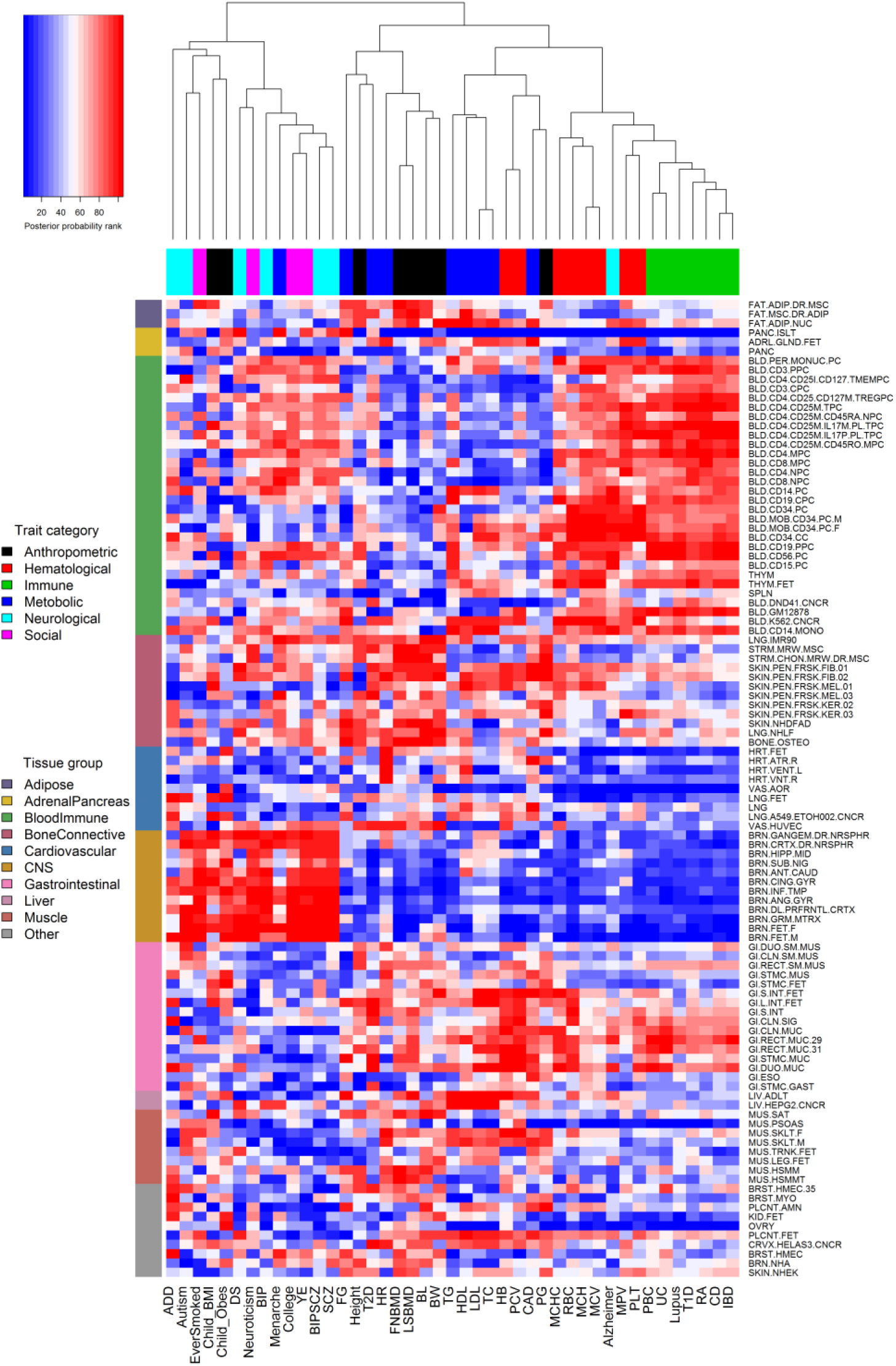
Heatmap displays the rank of 105 tissues (y-axis) in tenns of their relevance for each of the 43 GWAS traits (x-axis). Traits are organized by hierarchical clustering. Tissues are organized into ten tissue groups.

**Figure 5.**
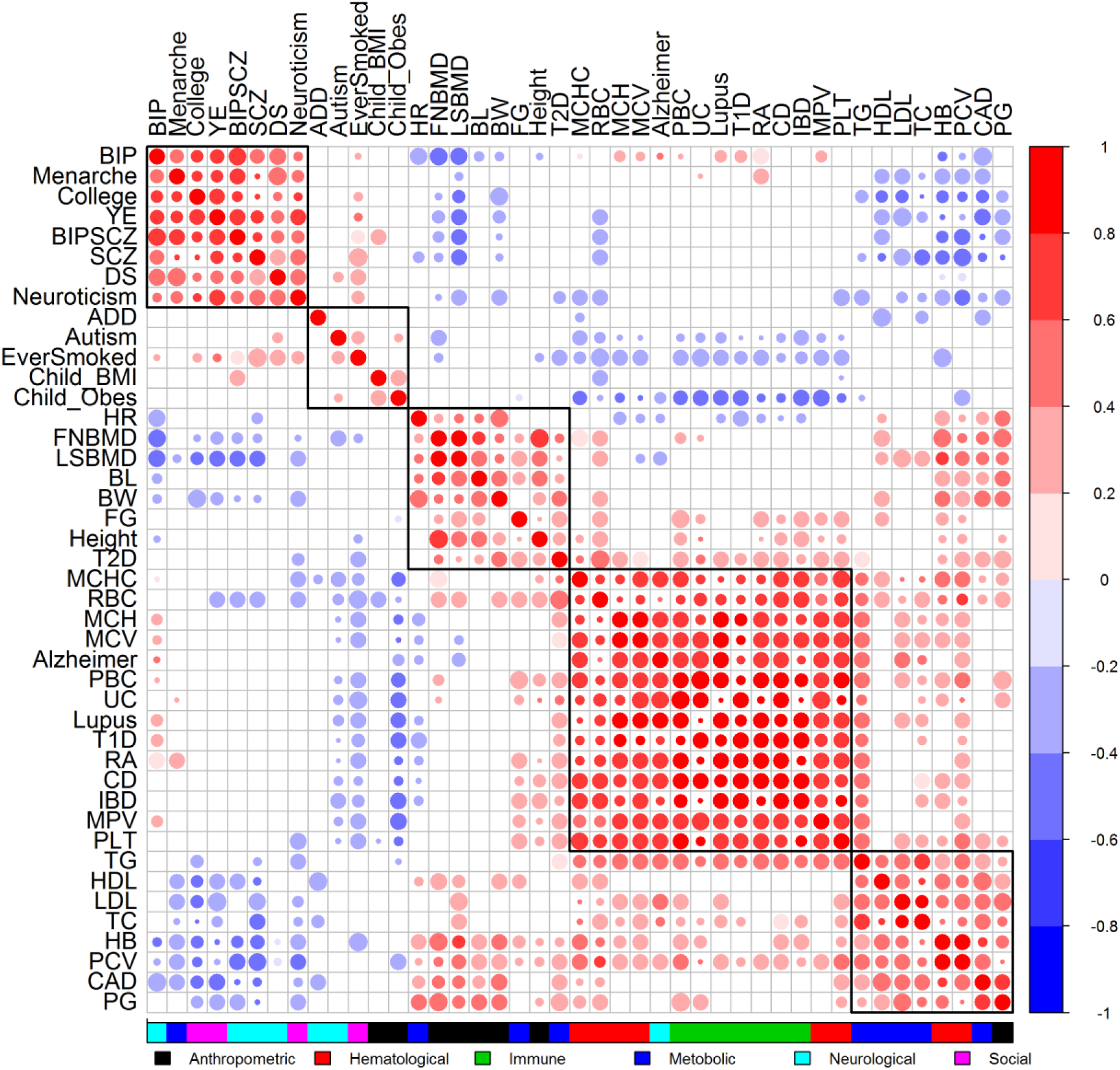
Correlation plot displays the relationship among 43 traits. Color of the circles in the correlation plot represents the sign and magnitude of the correlation coefficient (red: positive; blue: negative). No circle (while space) indicates no significant correlation (p > 0.05). Size of the circle indicates significance (large: p<0.001; median: p<0.01; small: p<0.05).

The first and second trait clusters contain psychiatric disorders (i.e. SCZ, BIP, Autism, DS, Neuroticism and ADD) and neurological related traits (e.g. College, YE, Menarche, Child_Obes, Child_BMI, and EverSmoked). For these two clusters, the CNS tissue tends to be identified as the most trait-relevant, consist with previous studies [16,27]. Among these traits, BIP, Menarche, College, YE, BIPSCZ, SCZ, DS, Neuroticism are highly correlated with each other and all belong to the first trait cluster; while ADD, Autism, EverSmoked, Child_BMI, Child_Obes are all grouped into the second trait cluster. Among the psychiatric trait pairs in the first two clusters, SCZ and BIP pair has the highest correlation (spearman correlation = 0.561; versus median/mean correlation = 0.098/0.167; BIPSCZ are excluded), suggesting that SCZ and BIP are more similar to each other in terms of tissue relevance footprint compared with the other trait pairs. The third trait cluster contains traits that are often related to several tissue groups. Specifically, the anthropometric traits (i.e. BL, BW, FNBMD, LNBMD and Height) are related to the bone, connective and gastrointestinal tissues; while the metabolic traits (i.e. FG, T2D and HR) are related to the gastrointestinal, liver and adipose tissues. The fourth trait cluster mainly contains two categories of traits that include immune diseases (e.g. RA, Lupus, T1D, UC, PBC, CD and IBD) and hematological traits (e.g. MCHC, MCH, MCV, PLT, RBC and MPV). Both these two categories are related to blood immune tissues. However, the fourth cluster also contains Alzheimer’s disease. The classification of Alzheimer’s disease in the fourth cluster rather than in the first two clusters presumably reflects the close relationship of the disease to both BloodImmune and CNS [28,74]. Finally, the fifth trait cluster mainly contains metabolic traits (TC, TG, LDL, HDL and CAD) that are related to gastrointestinal and blood immune tissues. Note that traits from the last three clusters tend to have positive correlations among each other, while have negative correlations with traits from the first two clusters (Figure 5).

### Using Trait-relevant Tissue Results in More Powerful SNP Set Tests

Finally, we explored the use of annotation coefficient estimates from the inferred trait-relevant tissues to construct SNP set tests in a separate data, the Wellcome trust cast control consortium (WTCCC) study. WTCCC contains the six common diseases that include T1D, T2D, CD, BIP, RA and CAD. We focused on a set of 5,588 genes and used 153,813 SNPs inside these genes to perform SNP set test using SKAT [49–51] (details in Materials and Methods). As in simulations, we considered three different SNP weights for SKAT test: (1) SNP weights constructed by the multivariate analysis approaches of SMART (i.e. ***HC*** and ***HB***); (2) SNP weights constructed by the univariate maximal statistics approach (***HCuMax, HBuMaxLDSC*** and ***HBuMax***); and (3) equal SNP weights (***EqualWeight***). We apply different weights to each of the six diseases. We first display the quantile-quantile (QQ) plot of −log10 p-values from SKAT in Figure 6 (for CD) and Figure S6 (for T1D, T2D, BIP, RA and CAD), which, consistent with simulations, suggests proper control of type I error. In the analysis, different approaches identified a different number of associated genes that pass the Bonferroni corrected genome-wide significance threshold (8.95×10^−6^), and these numbers range from 12–15 (the union of them contains 17 genes). These genes are associated with either CD, RA, T1D or T2D, and have all been validated either in the original WTCCC study or in other GWASs of the same trait (Table 1). Consistent with simulations, we found that SNP set tests using weights constructed from the trait-relevant tissue achieves higher power compared with using equal weights. For example, the ***HC*** approach or the ***HB*** approach identified 15 genes, 3 more than that identified using equal weights (Table 1). In particular, the ***HC*** approach identified ***C1orf141*** [77] and ***BSN*** [78] for CD as well as ***FTO*** [79] for T2D, and neither of these were identified by the equal weights approach. While the HB approach identified ***BSN*** [78] and ***SLC22A5*** [78] for CD as well as ***FTO*** [79] for T2D, and neither of these were identified by the equal weights approach. Finally, within each annotation set, using multiple annotations identified slightly more genes than using one annotation at a time (i.e. ***HB*** vs. ***HBuMax*** or ***HBuMaxLDSC*** and ***HC*** vs. ***HCuMax***), again consistent with the simulation results.

**Table 1.**
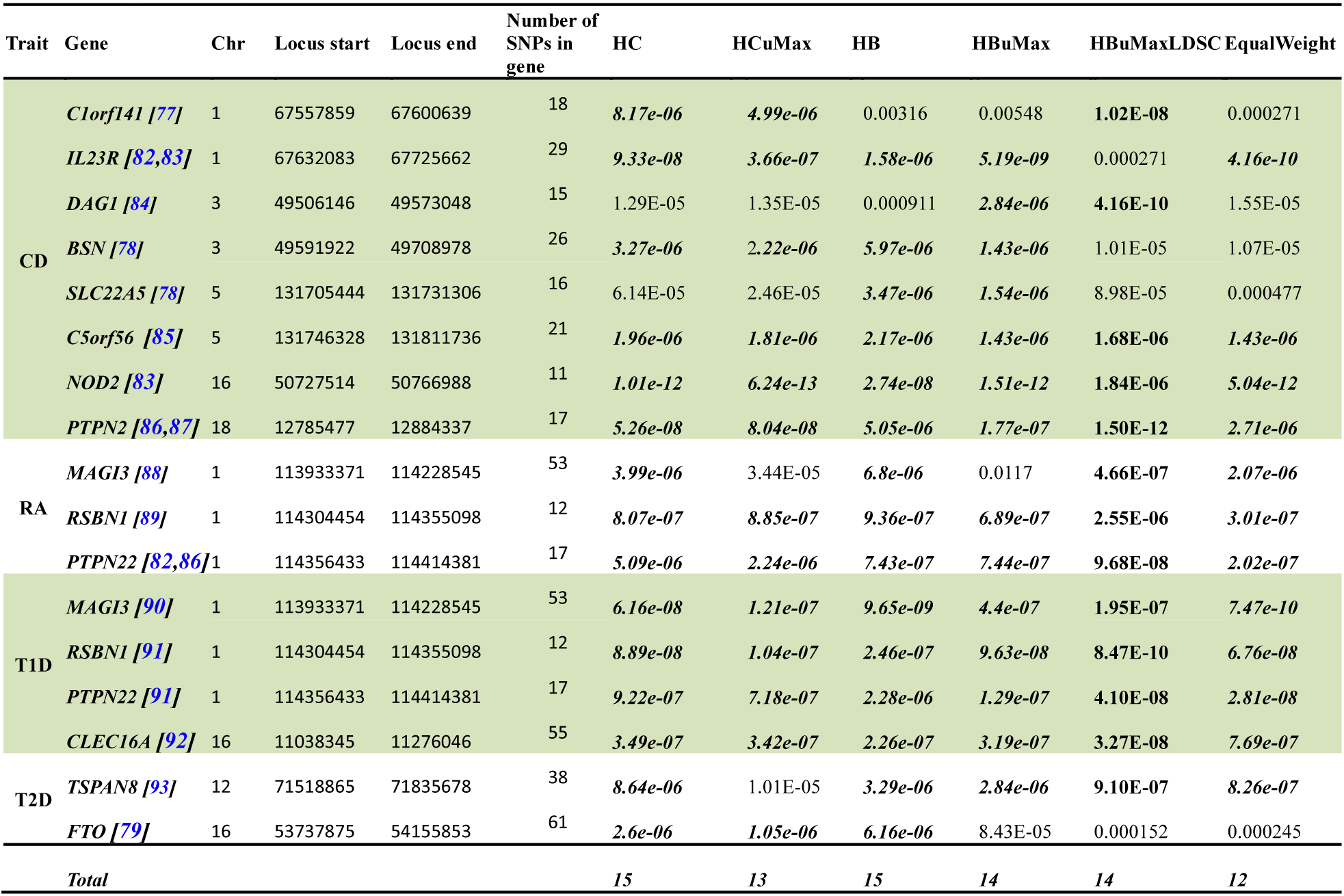
Association results for SNP set tests in WTCCC using different SNP weights. Results are shown for 17 genes identified to be significant by at least one SNP weighting option in four dieseases from the WTCCC data (CD, RA, T1D and T2D). All these genes have been previously identified to be associated with the corresponding trait (cited references). Approaches that yield a p-value passing the genome-wide significance threshold (8.95×10^−6^) are highlighted in bold.

**Figure 6.**
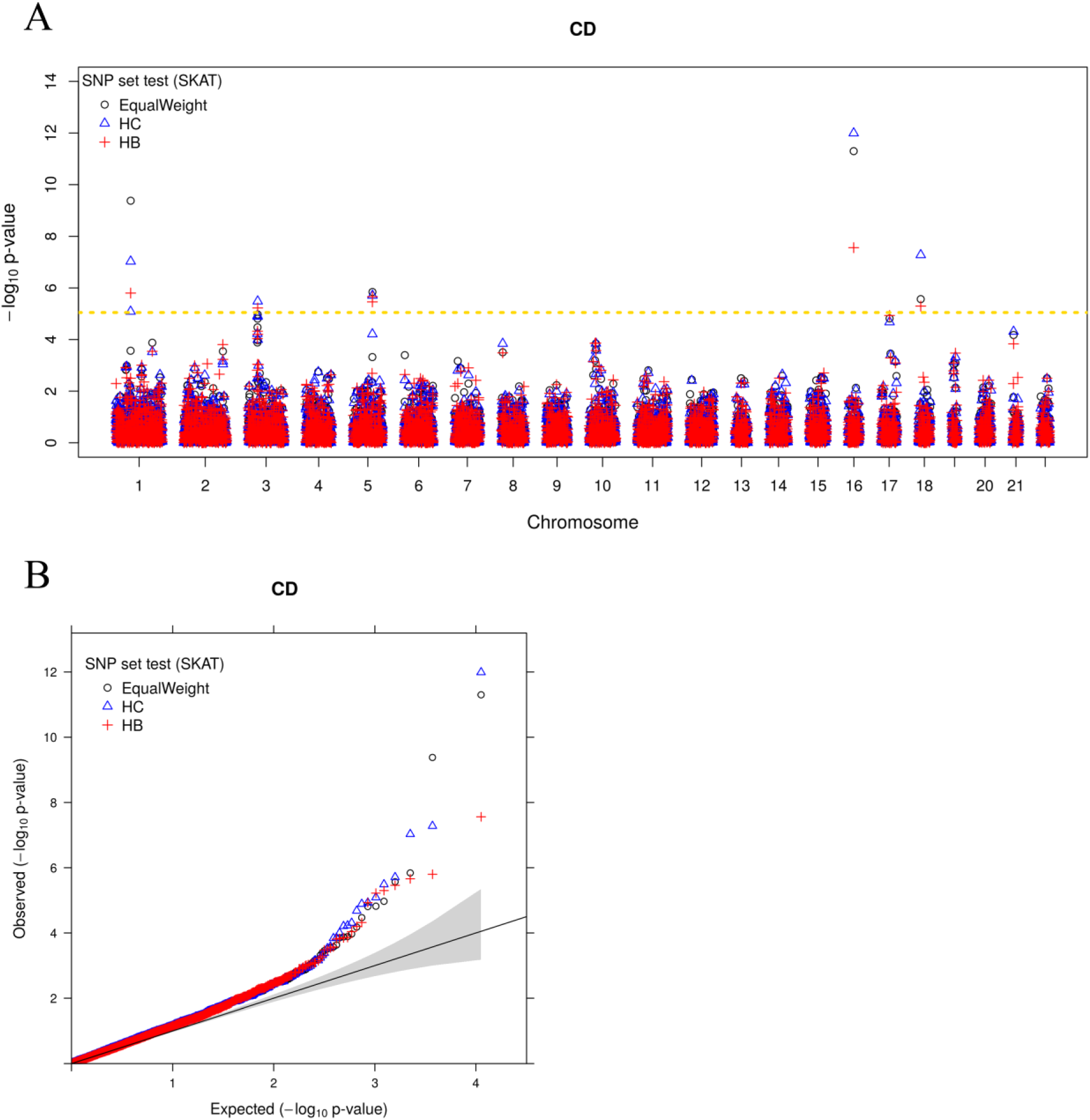
SNP set test results on Crohn’s disease (CD) using different SNP weights. (A) Manhattan plot shows association signal across genes (x-axis) detected by SNP set tests using three different sets of SNP weights. EqualWeight (black): equal SNP weights. HC (red): SNP weights constructed using the estimated coefficient parameters for continuous histone mark based annotations in the GWAS consortium study. HB (green): SNP weights constructed using the estimated coefficient parameters for binary histone mark based annotations in the GWAS consortium study. The gold dashed line represents genome-wide significance threshold (8.95×10^−6^). (B) The same results are displayed with QQ plot of −log10 p-values. Grey shaded area represents the 95% point-wise confidence interval.

## Discussion

We have presented a simple modification of the commonly used linear mixed model to integrate multiple SNP annotations with GWAS traits to facilitate the identification of trait-relevant tissues. We have described an accompanying GEE based parameter inference algorithm that makes use of summary statistics and naturally accounts for genetic correlation due to linkage disequilibrium. We have shown how the task of identifying trait-relevant tissues can be formulated into a classification problem and how mixture modeling can facilitate the inference of trait-relevant tissues in the presence of annotation correlation across tissues. We have also illustrated how the link between our extended linear mixed model and the commonly used SKAT can enable more powerful SNP set tests in new association studies. With both simulations and an in-depth analysis of 43 GWAS traits, we have illustrated the benefits of modeling multiple annotations jointly.

Our approach complements several recently developed methods that aim to derive a single, interpretable synthetic annotation by combining information of multiple annotations in a tissue specific fashion [17,20,27,28]. Most of these methods rely on multiple annotation information and use clustering algorithms to cluster SNPs into two categories. The posterior probability of SNPs in the category with the smaller number of SNPs thus becomes a synthetic annotation and is often interpreted as the posterior probability of being a “causal” or “functional” SNP. While these synthetic annotations have the benefits of simplicity and potential interpretability, they often have the drawback of being derived without taking into account the GWAS trait of interest. Arguably, functions of genetic variants depend on traits and clustering SNPs without considering the trait of interest may be suboptimal. Our approach complements these previous methods in that we effectively derive a synthetic annotation by taking GWAS traits into account. In particular, our method can be viewed as a supervised approach to combine multiple annotations into a single annotation, where the single annotation is represented as a weighted summation of the multiple annotations, with the weights being the estimated annotation coefficients inferred directly using the GWAS trait. Therefore, the approach we develop effectively takes the trait of interest into account. Certainly, both our approach and these previous approaches make a key modeling assumption that multiple annotations in the trait-relevant tissue are more relevant to SNP effect sizes or causality as compared with annotations in trait-irrelevant tissues. While it is a reasonable assumption for histone occupancy based annotations we examine here, this assumption may not hold well for certain annotations and for certain complex traits. For example, it is possible the classification of trait-relevant tissue depends on what annotation one examines: the SNP effect sizes can be predicted well by using one annotation in one tissue, or by using another annotation in a different tissue. To briefly explore the utility of our method in the case of multiple trait-relevant tissues, we performed a simulation study that is similar to the polygenic scenario presented in the results section but with two trait-relevant tissues: we used one annotation from one tissue and another annotation from another tissue to simulate SNP effect sizes. In this setting, as one might expect, the difference between the multivariate approach and the univariate approach is small (Figure S7). Therefore, developing method for the case of multiple trait-relevant tissues is an interesting future direction.

We rely on a polygenic model to evaluate the contribution of annotations to SNP effect sizes and infer trait-relevant tissues. Our polygenic model assumes that all SNP effect sizes are non-zero and follow a normal distribution with SNP-specific variance that is a function of multiple annotations. Therefore, our approach is different from several previous approaches that rely on a sparse model to evaluate the contribution of annotations to SNP causality [11,12,27,28,40,53]. While sparse models can be used to directly link annotation coefficients to the causality of SNPs, they often encounter severe computational burdens due to linkage disequilibrium among SNPs. For example, the recent sparse model bfGWAS [40] has to divide genome into thousands of approximately independent blocks and perform analysis within each block separately; and even with such a simplified algorithm it can take a sparse model days to analyze a GWAS data with tens of thousands of individuals and millions of SNPs. In contrast, a key advantage of our polygenic model and its GEE based inference algorithm is their ability to properly account for linkage disequilibrium while being computationally trackable. Indeed, it only take about 20 minutes to analyze each of trait-tissue pair in our real data application with hundreds of thousands of individuals and millions of SNPs. Certainly, the polygenic modeling assumption that all SNPs have non-zero effects may not be realistic for certain traits and developing both realistic and computationally efficient methods is an important future direction.

We have focused primarily on using tissue-specific annotations based on histone occupancy from the ENCODE and ROADMAP projects. Other tissue-specific annotations are nowadays readily available. For example, the GTEx project measures expression quantitative trait loci (eQTL) information in 53 tissues, many of which overlap with that in ROADMAP. Our method can easily incorporate multiple annotations from different data sources and include both eQTL annotations from GTEx and histone annotations from ROADMAP, though caution needs to be made to account for accuracy difference in these eQTL annotations from different tissues due to sample size variation. In any case, jointly analyzing multiple sources of annotations will likely improve power further in identifying trait-relevant tissues in the future.

In the real data application, we have attempted to infer fine-scale trait-tissue relevance by using 105 tissues instead of the 10 tissue groups using the ***HB*** approach. The inferred top-ranking tissue types from the ***HB*** approach for each of the 43 GWAS traits are listed in Supplementary Table S5. As expected, most of these top-ranking tissues belong to the top-ranking tissue group (median = 70.5% across traits), suggesting relatively stable inference results whether tissues or tissue groups were used in the analysis. For example, all of the top-ranking tissues (with PP > 0.5) for ever smoked and YE belong to the CNS tissue group, and 28 of the 39 top-ranking tissues for CD belong to the blood immune tissue group. We have attempted to further quantify the tissue-level relevance results by comparing them to the corresponding PubMed search results, as we have done in the main text for the tissue group analysis. However, we found that PubMed search results are unable to distinguish fine-scale tissue types for most traits. Therefore, we had to rely on prior biology knowledge obtained in various other studies to validate our tissue relevance analysis. In many cases, the top-ranking tissue fits our prior expectation. However, we also acknowledge that identifying relevant tissues from >100 tissue types is indeed a challenging task. Specifically, for 34 out of 43 traits, the PPs for more than half of the tissues within the corresponding top-ranking tissue group are greater than 0.5, suggesting that it is often difficult to identify a single trait-relevant tissue within the tissue group. Alternative approaches to explore fine-scale trait-tissue relevance have been suggested before. For example, a two-step analysis procedure was proposed to first identify trait-relevant tissue group and then identify trait-relevant tissue within the tissue group [80]. In addition, using synthetic annotations generated from Genoskyline [28] or FUN-LDA [81] could be particularly useful for identifying fine scale trait-relevant tissues. Our method can be easily adapted to incorporate a two-step analysis procedure and/or accommodate synthetic annotations, and has the potential to yield better trait-tissue relevance resolution in the future.

Finally, while we have mainly focused on inferring trait-relevant tissues, we have also explored the feasibility of using inferred trait-relevant tissues and the estimated annotation coefficients to enable more powerful SNP set test in future GWASs. In practice, multiple annotation sets can be used to construct SNP set tests (e.g. HC and HB annotations sets as used in our real data application). It is often difficult *a priori* to determine which annotation set would yield the best results. Therefore, we recommend analyzing all of these annotation sets separately and choose the one that yields the highest power, as we have done in the real data application. In addition, sometimes the trait of interest may have multiple relevant tissues. In this case, we can apply the PPs from the identified trait-relevant tissues (with PP>0.5) to weight the corresponding estimated annotation coefficients from these tissues to form a set of weighted annotation coefficients, in line with the Bayesian model averaging idea. Finally, while incorporating annotation information does increase SNP set test power, we also found that such power improvement in realistic settings depends on traits and can be variable. The variability in power improvement of our method is consistent with many previous studies that have shown similar variability in power improvement by integrating SNP annotations into single SNP association tests [11,40]. However, increasing sample size and the development of better SNP annotations will likely facilitate the adaption of various annotation integration methods in the near future.

## Supporting Information Legends

**Figure S1** Simulation results for comparing using multiple annotations versus using a single annotation. Power to detect trait-relevant tissues by different approaches in various settings at a fixed FDR of 0.05 (A), 0.1(B), or 0.2 (B). x-axis shows the values of the two annotation coefficients used in the simulations. Settings where at least one annotation coefficient is zero are shaded in grey. The setting where the annotation coefficients equal to the median estimates from real data (i.e. α = (0.1, 0.05)) is shaded in gold.

**Figure S2** Per-block PVE of the ten causal blocks for the second set simulations. (A) The cases of one informative annotations at **α_0_ = 0. 5** and (**α_1_, α_2_**) = (**0. 4,0**); (B) The cases of two informative annotations at **α_0_** = **0. 5** and (**α_1_, α_2_**) = (**0.4, 0.4**).The bar indicates the standard error across simulation replicates.

**Figure S3** Various annotation characteristics influence the power in identifying trait-relevant tissues in simulations. Methods for comparison include SMART (red), UniMax (green), and UniMaxLDSC (blue). Area under the curve (AUC) is used to measure method performance. (A) Power to identify trait-relevant tissue generally increases with increasingly large annotation coefficients when the two coefficients have the same sign. (B) Power also increases with increasingly large annotation coefficients when the two coefficients have the opposite sign. (C) Power is relatively stable with the genome coverage of the two annotations varied from 4%, 8% to 12%.

**Figure S4** Various factors influence the power of SNP set analysis in simulations. Left columns (A, D, G): annotation coefficients are fixed to be (1, 1) while the number of causal blocks changes from 5, 10, 20 to 50. Middle columns (B, E, H): the number of causal blocks is fixed to be 10 while the annotation coefficients change from (0.01, 0.01), (0.3, 0.3), (0.6, 0.6) to (1, 1). Right columns (C, F, I): per-block PVE are approximately fixed while the number of causal blocks and annotation coefficients vary. Top rows (A, B, C) show the average proportion of phenotype variance explained (PVE) by non-causal or causal blocks. Middle rows (D, E, F) show the fold enrichment. Bottom rows (G, H, I) show SNP set analysis power for various methods.

**Figure S5** Posterior probabilities of 10 tissue groups for being relevant to each of the 43 GWAS traits by HC. The gold dashed line represents a horizontal line at 0.5.

**Figure S6** Manhattan and QQ plots display the SNP set test results for five common diseases in WTCCC using different SNP weights. Results are shown for rheumatoid arthritis (RA; A), cardiovascular disease (CAD; B), bipolar disease (BIP; C), type II diabetes (T2D; D), and type I diabetes (T1D; E). For comparison, association results based on univariate SNP tests are also shown in F-K. EqualWeight (black): equal SNP weights. *HC* (blue): SNP weights constructed using the estimated coefficient parameters for continuous histone mark based annotations in a GWAS consortium study. *HB* (red): SNP weights constructed using the estimated coefficient parameters for binary histone mark based annotations in a GWAS consortium study. For Manhattan plots, gold dashed lines represent genome-wide significance thresholds: 0.05/153,813 for univariate tests and 0.05/5,588 for SNP set tests. For QQ plots, grey shaded area represents the 95% point-wise confidence interval.

**Figure S7** Simulation results for SKAT. As the second set of the simulations, 10,000 individuals and 10,000 SNPs were selected from GERA study. The SNPs were divided into 100 blocks with 100 SNPs in each block. Two histone marks were simulated for 40% SNPs in the causal blocks of the trait-relevant tissue. Left panel (A, D, G): Fix the annotation effect (1, 1) and change the number of causal blocks; Middle panel (B, E, H): Fix the number of causal blocks 10, and the change the annotation effects (0.01, 0.01), (0.3, 0.3), (0.6, 0.6) and (1, 1); Right panel (C, F, I): Fix the per-block PVE, and change the number of causal blocks and annotation effects.

**Table S1** Information for the tissue-specific SNP annotations obtained based on histone occupancy data in the ENCODE and Roadmap projects. The table lists ID, tissue group, epigenome name and mnemonic, tissue types, and genome-wide percentage of mark occupancy for the four histone marks (H3K27me3, H3K36m3, H3K4me1, H3K4me3).

**Table S2** Information for the summary statistics of 43 traits from 29 GWAS studies. The table lists the phenotype name, category, abbreviation, number of individuals, reference, and downloaded websites for each of the 43 traits.

**Table S3** The most relevant tissue group for each trait determined by various methods. Parenthesis shows either the proportion of existing publications for the tissue group (for PubMed search), the tissue group level posterior probability for other approaches (HC, HCuMax, HB, HBuMax, Chmm, cHMMuMax), or the tissue group level Wald statistics for HBuMaxLDSC.

**Table S4** PubMed search results show the number (in parenthesis) and the normalized proportion of publications on each pair of tissue group (columns) and trait (rows). The PubMed search was performed on June 23, 2017. For each GWAS trait, the tissue group with the largest proportion of existing publications are highlighted in red. The proportion values in each row sum to one.

**Table S5** The posterior probability of 105 tissues with four histone marks for 43 complex traits inferred by SMART.

